# Open Design 3D-Printable Adjustable Micropipette that meets ISO Standard for Accuracy

**DOI:** 10.1101/109231

**Authors:** Martin D. Brennan, Fahad F. Bokhari, David T. Eddington

**Author notes:** Correspondence and requests for materials should be addressed to David T. Eddington.

## Abstract

Scientific communities are drawn to the open source model as an increasingly utilitarian method to produce and share work. Initially used as a means to develop freely available software, open source projects have been applied to hardware including scientific tools. Increasing convenience of 3D printing has fueled the proliferation of open labware projects aiming to develop and share designs for scientific tools that can be produced in-house as cheap alternatives to commercial products. We present our design of a micropipette that is assembled from 3D-printable parts and some hardware that works by actuating a disposable syringe to a user adjustable limit. Graduations on the syringe are used to accurately adjust the set point to the desired volume. Our open design printed micropipette is assessed in comparison to a commercial pipette and meets ISO 8655 standards.

## Introduction

The open source development model, initially applied to software, is thriving in the development of open source scientific equipment due in part to increasing access of 3D printing^1,2^. Additive manufacturing methods have existed for decades although the recent availability of inexpensive desktop printers^3,4^ have made it feasible for consumers to design and print prototypes and even functional parts, and consumer goods^5,6^. Proliferation of free CAD software^7–10^ and design sharing sites^11–14^ have also supported the growth and popularity of open designed parts and projects. Open design 3D-printable lab equipment is an attractive idea because, like open source software, it allows free access to technology that is otherwise inaccessible due to proprietary and/or financial barriers. Open design tools create opportunity for scientists and educational programs in remote or resource limited areas to participate with inexpensive and easy to make tools^15–20^. Open source development also enables the development of custom solutions to meet unique applications not met by commercial products that are shared freely and are user modifiable^5,21–26^. Some advanced, noteworthy, open source scientific equipment include a PCR device^27^, a tissue scaffold printer^28^, and a two-photon microscope^29^ although simple tools have potential to be impactful as they can serve a wider community.

Some simple and clever printable parts that have emerged are ones that give a new function to a ubiquitous existing device, such as drill bit attachment designed to hold centrifuge tubes, allowing a dremel to be used as a centrifuge^30^. Although this may make a rather crude centrifuge it may be an adequate solution for a fraction of the cost of a commercial centrifuge. Other examples of open source research tools that utilize 3D printed parts include optics equipment^31^, microscopes^32,33^, syringe pumps^34,35^, reactionware^36–39^, microfluidics^40,41^ and the list continues to grow^42^.

One example of a everyday scientific tool that provides opportunity for an open design solution is the micropipette. An open design micropipette that can be made cheaply affords more options for labs and educational settings. Micropipettes are an indispensable tool used routinely in lab tasks and can easily cost $1000 USD for a set. Often a lab will require several sets each for a dedicated task. Some pipettes may even re-calibrated for use with liquids of different properties.

Air displacement pipettes use a piston operating principal to draw liquid into the pipette^43^. In a typical commercial pipette the piston is made to be gas-tight with a gasketed plunger inside a smooth barrel. Consumer grade fused deposition modeling (FDM) printers are unable to build a smooth surface do to the formation of ridges that occur as each layer is deposited^44^. The ridges formed by FDM make it impractical to form a gas-tight seal between moving parts, even with a gasket. Existing printable open-design micropipettes get around this limitation by stretching a membrane over one end of a printed tube, which when pressed causes the displacement. The displacement membrane can be made from any elastic material such as a latex glove. A few open design micropipettes exist including a popular one which, in addition to the printed parts, uses parts scavenged from a retractable pen^45^. Because there is no built in feature such as a readout for the user to set the displacement to a desired volume, this design requires the user to validate the volumes dispensed with a high precision scale. Without verification with a scale the volumes dispensed can only be estimated based on calculations of the deflection of the membrane which is not a practical protocol.

We submit a new design whose major strength is the ability to adjust to any volume aided by the the built in scale. Our open design 3D-printable micropipette works by actuating a disposable syringe to a user adjustable set point. This allows the user to set the pipette to a volume by reading the graduations on the syringe barrel. We have also designed an adjusted graduation scale, to be used in place of the printed on scale, that corrects for the compressibility of air which allows the pipette to be set accurately. Additionally our pipette offers a simplified assembly requiring no glue, tape, or permanent connections.

## Results and Discussion

Validation testing preformed with the existing graduations printed on the syringes did not meet the ISO standards. According to ISO 8655 for a pipette with the maximum nominal volume of 1000 *µ*L the systematic error cannot exceed 8 *µ*L and the random error cannot exceed 3 *µ*L. For the maximum nominal volume of 300 *µ*L the systemic error cannot exceed 4 *µ*L and the random error cannot exceed 1.5 *µ*L. The commercial pipettes met these standards handily, but in our initial tests with our printed pipette we noticed large negative systematic error suggesting that we were missing a biasing factor. This error was exaggerated while transferring larger volumes. After further investigation we realized that the graduations are intended to measure incompressible fluids within the syringe while we were using them to measure air while under vacuum. Because the syringe graduations were used to measure the air under vacuum it could not be expected to serve as a 1:1 volume ratio of water that was pulled into the tip because under vacuum the air had expanded. This explains the negative systematic error we were experiencing. The water in the tip pulls on the air due to gravity which causes it to expand slightly but enough to make measuring with the graduation in this way inaccurate. Our solution was to replace the built-in scale with a scale that accounted for the expansion. Using the data from our initial testing we calculated a factor for the expansion and used that to make the new scale (See SI Fig. 1 and SI Fig. 2). With the new scale taped on the syringe our validation testing met the ISO 8655 standard. Replacing the scale with our scale that accounts for the expansion corrected this effect. Our printed pipette with the adjusted scale meets ISO standards for accuracy and precision. (Table 3 and 4).

**Figure 1:**
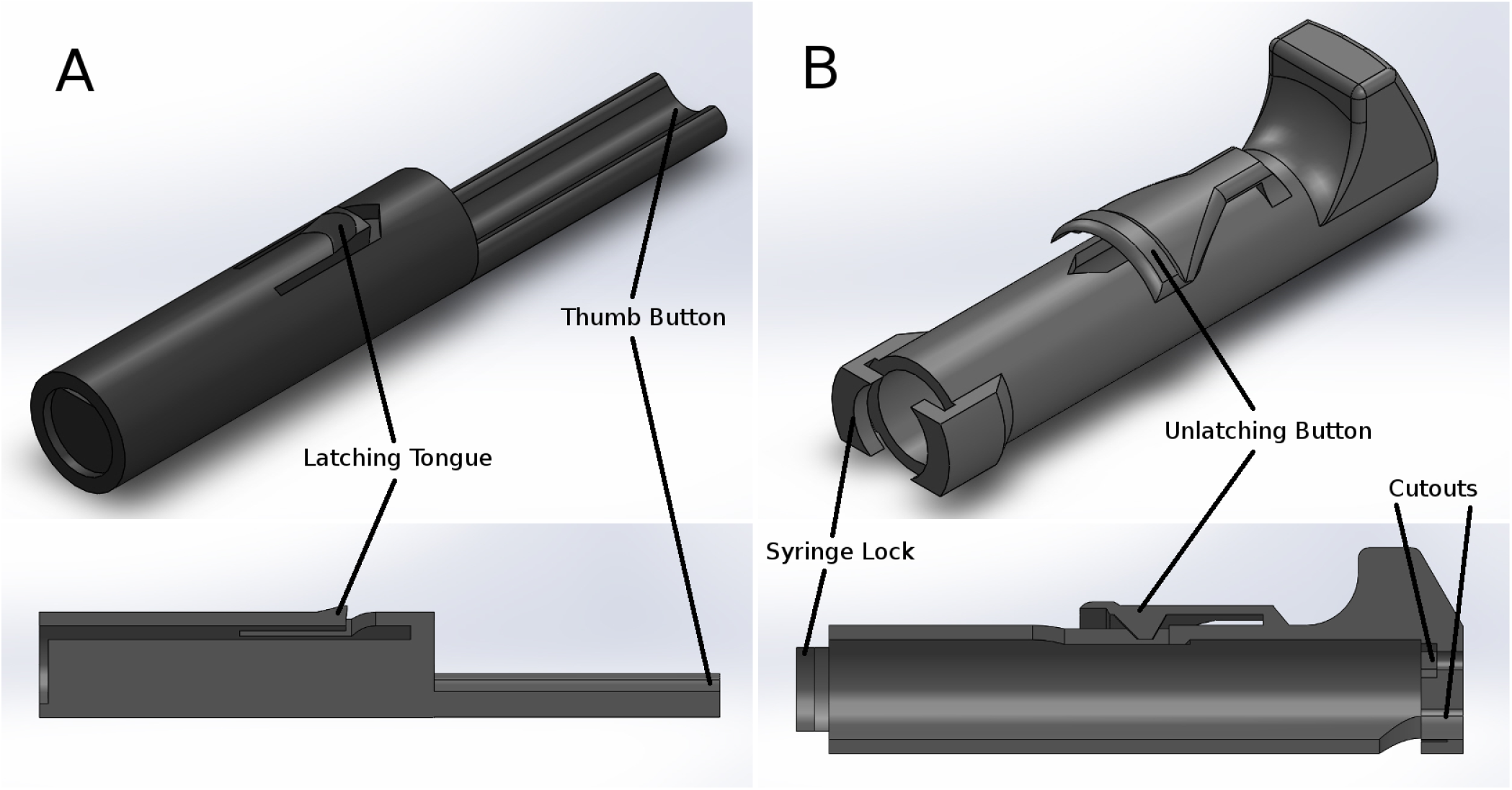
CAD renderings of printable parts and cross-sections. (A) The printed plunger part is a shaft pushes on the syringe plunger and slides in the printed body part. The printed plunger has a latching tongue and button which interfaces with the printed body part. (B) The printed body part holds the syringe and interfaces with the printed plunger part. The body part features the unlatching button and slots to hold the syringe in place by its flanges. The body part also has two cutouts in the top for the plunger button and for the hex nut and bolt.

**Figure 2:**
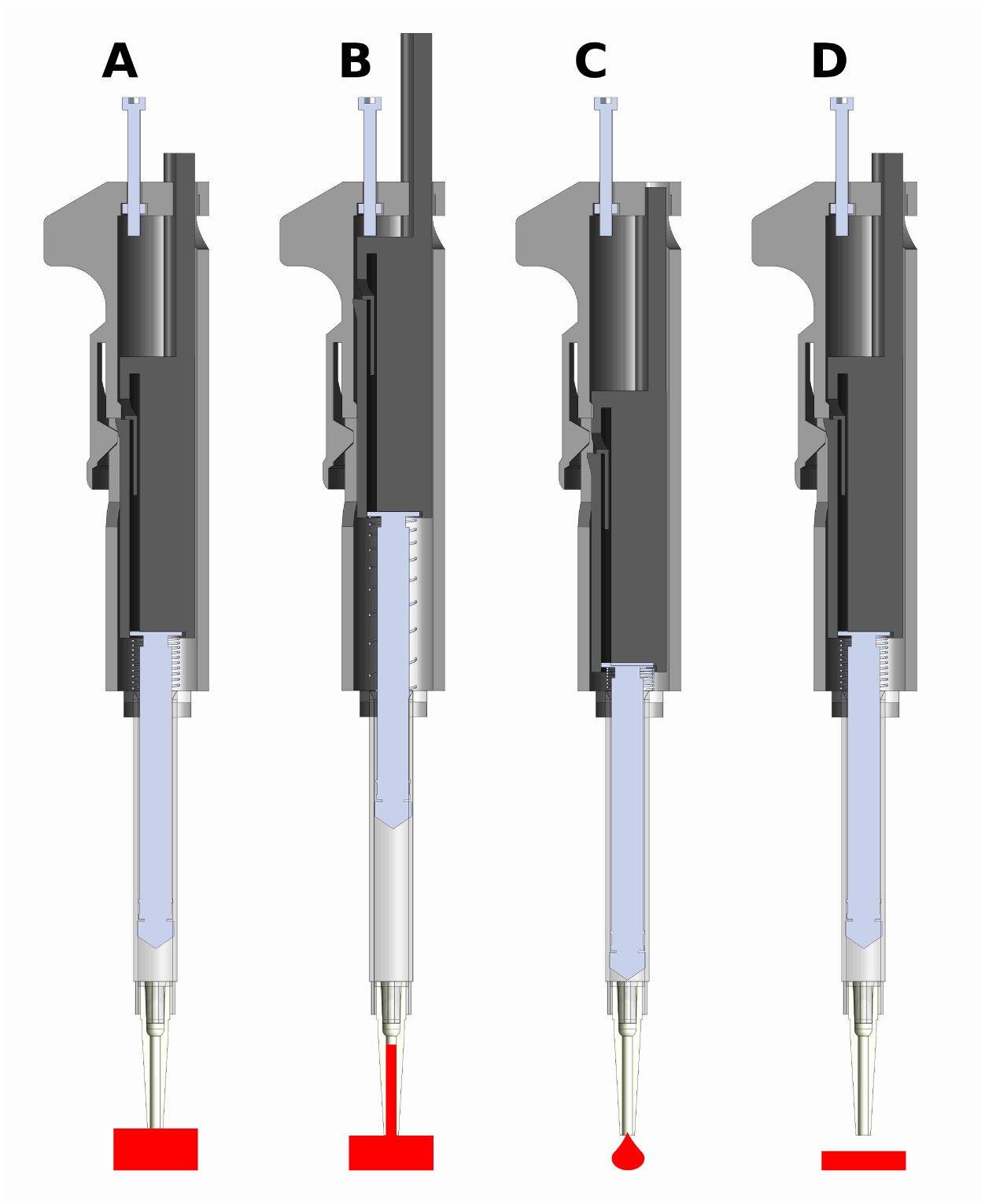
CAD renderings of assembled pipette and function. The pipette actuates the syringe to three positions. **(A) The latched position.** When the plunger is pressed the pipette locks at this position. The tip is then placed in a liquid and the unlatching button is pressed to release the pipette back to the set position (B), drawing in liquid. **(B) The set position.** The position of the screw determines the total displacement that the plunger moves. The pipette is spring loaded to return to this position. **(C) The blow-out position.** The fluid is transferred by pressing the plunger past the latched position to blow-out all the liquid. **(D) Return to the latched position.** The pipette returns to the latched position ready to preform another transfer.

**Figure 3:**
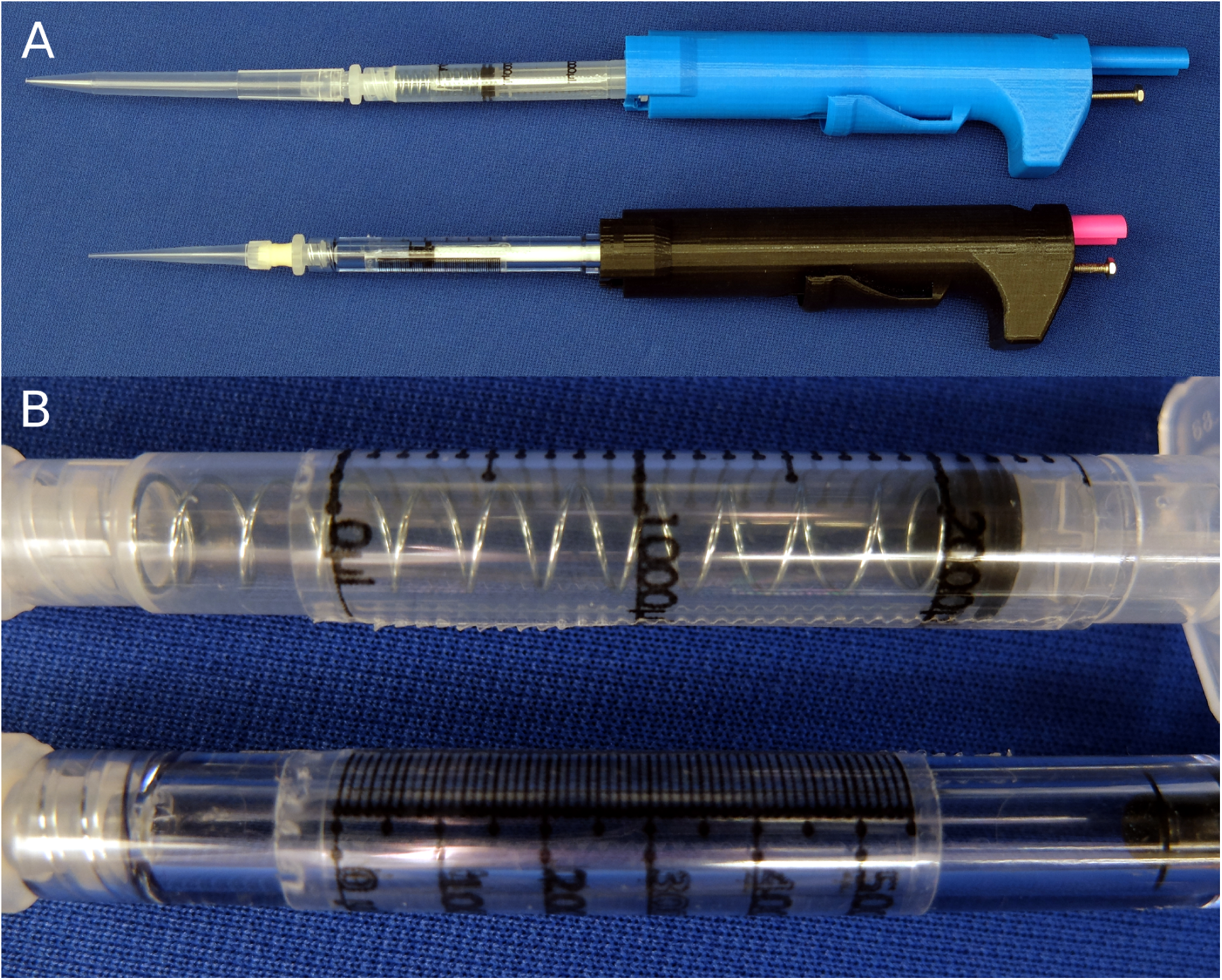
Photos of assembled pipettes. (A) Two assemblies of the pipette: the 100-1000 *µ*L configuration (above) and the 30-300 *µ*L configuration (below). (B) Close-up photo of the taped on scale for each of the syringes.

Our printed pipette improves on existing open design pipettes in several ways. Most significantly the user is able to adjust the syringe accurately without verifying the volume with a scale. The pipette can be adjusted to any discrete volume within range. In addition, the assembly requires no permanent connections using tape or glue which allows for re-configuration and easy replacement of parts. Conveniently, our design can also reach to the bottom of a 15 mL conical tube allowing tasks such as aspirating supernatant fluid from a cell pellet. Unlike existing designs our design does not use a membrane that may wear out and require tedious replacement. The only major limitation of this design compared to the biropipette is the option for a pipette tip ejector. The biropipette also uses arguably easier to source parts (a pen VS a syringe) relative to what is required from our design. The biropipette also was developed in OpenSCAD which is free and open source CAD software where our design was created in Solidworks a proprietary software.

This design relies heavily on the syringe, a specialized part, for the core function as well as accuracy. Fortunately, as disposable syringes are a regulated medical device they must meet high standards making them a reliable part for this design. As good as the performance of our printed pipette is, a commercial pipette should be expected to be more reliable and preferred for critical procedures. Adjusting our printed pipette to the desired volume requires more time and attention which would be tedious for procedures requiring many adjustments. On the other hand, due to the $6 cost to make, many printed pipettes could be assembled and pre-set, each for a planned transfer, rather then re-adjusting one commercial pipette for each step. The printable pipette can also be used as a disposable, sparing the commercial micropipette, in situations were cross-contamination and resterilization is required, or damage from volatile reagents is a concern. This pipette would also be ideal for high school and teaching labs or any resource limited program especially as part of a module about 3D printing.

The open design nature of this project encourages anyone to use and submit changes to improve and add features to this design. All of the working files and documentation are kept in a public repository ^46^. Future directions for this project include developing a tip ejection system and additional configurations for transferring volumes below 30 uL as well as increasing over all user friendliness, such as making the adjusting bolt easier to grip and improving ergonomics.

## Conclusion

We have presented an open design micropipette that uses 3D-printable parts in addition to a disposable syringe and a few easily sourced pieces of hardware. Our open design pipette is novel in that it allows the user to set the pipette to the desired volume without the need to calibrate or verify with a weighing scale. We validated the accuracy and precision of our pipette and developed a new graduation scale to correct for the use of the syringe for air-displacement measurements that meets the ISO standard for adjustable pipettes. Our design is free to use and modify to encourage further collaborative development.

## Methods

Our printed pipette is designed to actuate a 1 mL or 3 mL syringe to a user set displacement. The core of the design is two printed parts, the body and the plunger which are able to be printed on a consumer grade FDM printer (fig. 1 1), in this case a Makerbot Replicator. A 1 mL or 3 mL syringe twists to lock in the body part and is held into place by the syringe flanges. The 30-300 *µ*L configuration uses a 1 mL pipette and the 100-1000 *µ*L configuration uses a 3 mL pipette. The plunger part slides freely in the body part and actuates the syringe by pushing the thumb button (fig. 2 2). The pipette is spring loaded towards a set point which is adjustable by a set-screw. When the thumb button is pressed the system locks when it reaches the latched position, where it is ready to draw in fluid. The plunger is held in place with a latching button design which is released with the unlatching button drawing in fluid. The displacement is equal to the distance between the set position and the latched position. The pipette can also be pressed past the latched position to ‘blow-out’ the transferred fluid completely from the pipette tip. Additional materials required for assembly include two springs, a nut, a bolt and two washers. Attempts to make a printable luer lock adapter for tips was abandoned as the surface of printed parts is too rough to make an air tight seal with the luer or pipette tip. Instead, a combination of a barbed luer adapter and elastic tubing is used to attach the pipette tips. Our printed pipette mimics commercial pipettes design, function, and user operation, making it intuitive to use.

Our pipette uses the air-displacement method, where a vacuum is applied to a pocket of air to draw liquid into the pipette. As air is an compressible fluid, this pocket of air grows due to the weight of the liquid pulling on it. Due to this effect the graduations on the syringe are not accurate, as they are designed for measuring liquid within the syringe. At larger volumes this effect is more pronounced resulting in the volume measured being greater than the amount of liquid pulled into the syringe. We remedied this by creating a new scale to account for the expansion. The scale is printed on a transparency sheet and is taped onto the syringe for accurate measurements (See SI Fig. 1 and SI Fig. 2).

### Fabrication & Assembly

Two parts are printed (body.stl and plunger.stl) at normal resolution with rafts on a Makerbot Replicator. A small amount of paraffin wax is applied to the screw to prevent slop from causing the set point to drift after each actuation. The nut is sunk into the hex inset in the printed body part. The bolt is threaded in from the top of the body into the nut. Springs and washers are threaded onto the plunger of a 1 mL syringe for the 30-300 *µ*L configuration. Springs are placed inside a 3 mL syringe for the 100-1000 *µ*L configuration. The plunger part is inserted in the body and the syringe assembly is pushed in and locked from the syringe flanges to the body part to complete assembly (fig. 3 3). Parts and cost are listed in Table 1 and 2.

**Table 1:** Parts and cost for the 30-300 *µ*L pipette.

| Part | Unit Price | Source | Part number |
| --- | --- | --- | --- |
| Filament | \$1.63 | Makerbot | NA |
| 1 mL Syringe | \$0.15 | BD Biosciences | 309628 |
| Bolt | \$0.12 | McMaster-Carr | 91287A026 |
| Nut | \$0.01 | McMaster-Carr | 90591A121 |
| Spring (2) | \$4.14 | McMaster-Carr | 94125K542 |
| Washers (2) | \$0.16 | McMaster-Carr | 90107A012 |
| total | \$6.21 | | |

**Table 2:** Parts and cost for the 100-1000 *µ*L pipette.

| Part | Unit Price | Source | Part number |
| --- | --- | --- | --- |
| Filament | \$1.63 | Makerbot | NA |
| 3 mL Syringe | \$0.73 | BD Biosciences | 309657 |
| Bolt | \$0.12 | McMaster-Carr | 91287A026 |
| Nut | \$0.01 | McMaster-Carr | 90591A121 |
| Spring (2) | \$4.14 | McMaster-Carr | 94125K542 |
| total | \$6.63 | | |

**Table 3:**
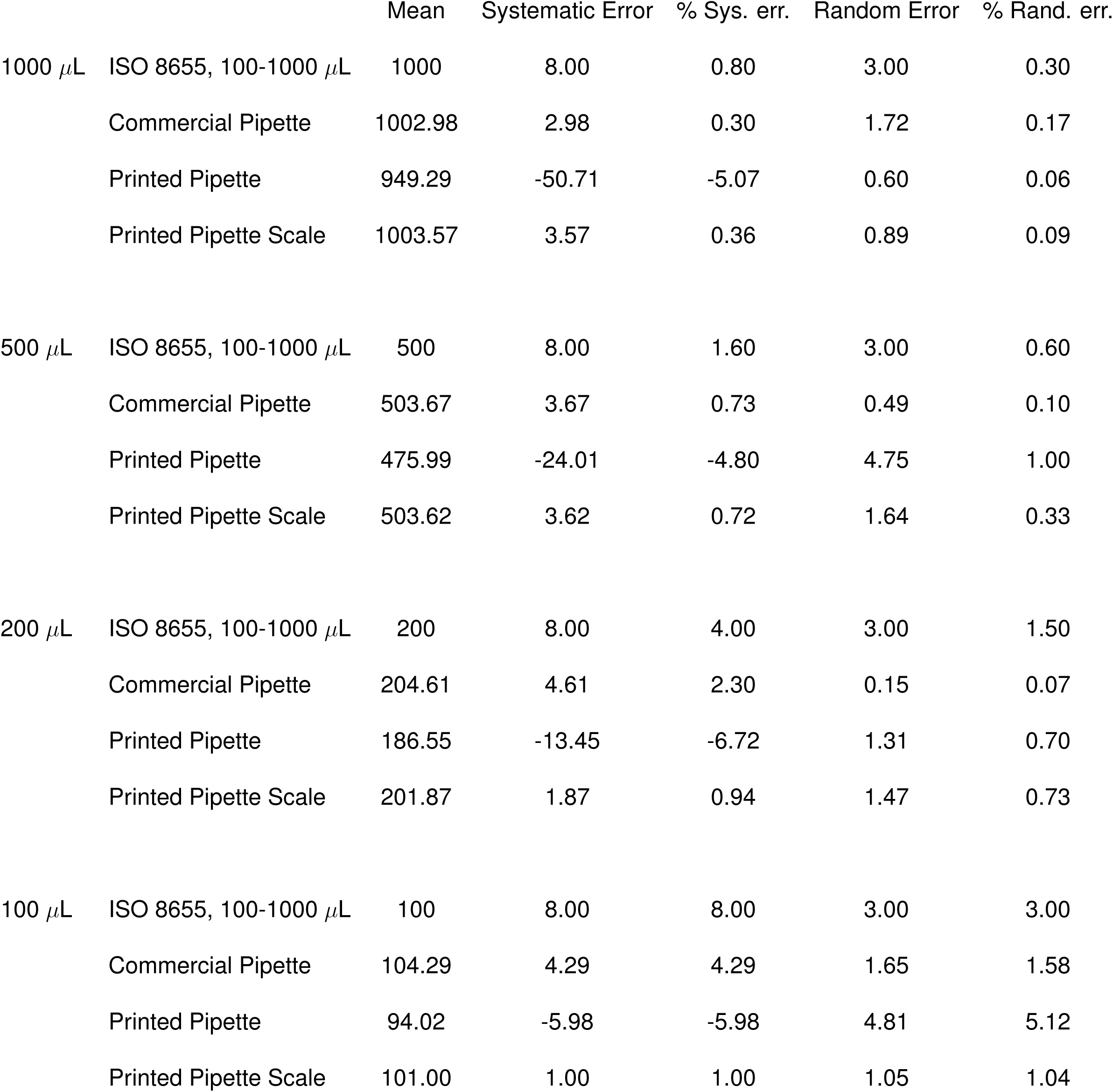
ISO 8655 for 100-1000 *µ*L comparing a commercial pipette with our printed pipette used with existing 3 mL syringe scale and an adjusted scale

**Table 4:** ISO 8655 for 30-300 *µ*L comparing a commercial pipette with our printed pipette used with existing 3 mL syringe scale and an adjusted scale

|  |  | Mean | Systematic Error | % Sys. err. | Random Error | % Rand. err. |
| --- | --- | --- | --- | --- | --- | --- |
| 300 $\mu\text{L}$ | ISO 8655, 30-300 $\mu\text{L}$ | 300 | 4.00 | 1.33 | 1.50 | 0.50 |
|  | Commercial Pipette | 301.19 | 1.19 | 0.40 | 0.53 | 0.18 |
|  | Printed Pipette | 286.91 | -13.09 | -4.36 | 0.42 | 0.15 |
|  | Printed Pipette Scale | 299.11 | -0.89 | -0.30 | 0.48 | 0.16 |
| 200 $\mu\text{L}$ | ISO 8655, 30-300 $\mu\text{L}$ | 200 | 4 | 2 | 1.5 | 0.75 |
|  | Commercial Pipette | 200.06 | 0.06 | 0.03 | 0.46 | 0.23 |
|  | Printed Pipette | 193.40 | -6.60 | -3.30 | 2.86 | 1.48 |
|  | Printed Pipette Scale | 200.57 | 0.57 | 0.28 | 0.86 | 0.43 |
| 50 $\mu\text{L}$ | ISO 8655, 30-300 $\mu\text{L}$ | 50 | 4 | 8 | 1.5 | 3 |
|  | Commercial Pipette | 49.02 | -0.98 | -1.96 | 0.10 | 0.20 |
|  | Printed Pipette | 49.62 | -0.38 | -0.76 | 1.26 | 2.53 |
|  | Printed Pipette Scale | 48.73 | -1.27 | -2.54 | 1.11 | 2.27 |
| 30 $\mu\text{L}$ | ISO 8655, 30-300 $\mu\text{L}$ | 30 | 4 | 13.3 | 1.5 | 5 |
|  | Commercial Pipette | 29.08 | -0.92 | -3.06 | 0.09 | 0.31 |
|  | Printed Pipette | 29.22 | -0.78 | -2.59 | 0.31 | 1.07 |
|  | Printed Pipette Scale | 27.78 | -2.22 | -7.41 | 1.37 | 4.93 |

### Validation

The printed pipettes accuracy and precision was characterized and compared to a commercial pipette as well as ISO 8655. The printed pipette was adjusted to the target volume by eye from the syringe graduations. Deionized water was transferred and measured with a scale. Five transfers were recorded and averaged to account for random variability. Data was taken for printed pipettes with existing syringe graduations as well as with the our adjusted scale. Data was taken with commercial pipettes of 30-300 *µ*L and 100-1000 *µ*L to compare to the printed pipette. Accuracy and precision are expressed as systematic error and random error, respectively, and are calculated according to ISO 8655 ^43^ (Table 3 and 4). The systematic error, or accuracy, is calculated according to the following equations where accuracy (*A*) is the difference of the mean volume 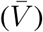 and the nominal volume (*V*_*o*_):

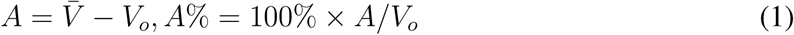

The precision or random error is the standard deviation (*s*) of the measurements and (*cv*) is the coefficient of variation:

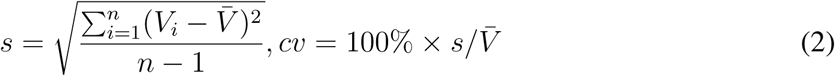

## Acknowledgements

We would like to acknowledge the UIC Bioengineering Senior Design team of 2014 for initial work on this project: Oluwafemi Aboloye, Pedro Hurtado, Rafael Romero, and Vivian Sandoval.

